# The TIRR–53BP1 axis controls PLK1 spatiotemporal regulation during mitosis

**DOI:** 10.64898/2026.06.29.735294

**Authors:** Aniruddha Sarkar, Shrabasti Roychoudhury, Katherine N. Choe, Neil T. Umbreit, H. Rudolf de Boer, Yizhou Joseph He, Bartlomiej Tomasik, Marcel A. T. M. Vugt, David Pellman, Dipanjan Chowdhury, Alexander Spektor

**Affiliations:** Division of Radiation and Genome Stability, Department of Radiation Oncology, Dana-Farber Cancer Institute, Harvard Medical School, Boston, MA 02215, USA; Department of Cell Biology, Blavatnik Institute, Harvard Medical School, Boston, MA, USA; Department of Pediatric Oncology, Dana-Farber Cancer Institute, Boston, MA, USA; Single-Cell Sequencing Program, Dana-Farber Cancer Institute, Boston, MA, USA; Howard Hughes Medical Institute, Chevy Chase, MD, USA; Broad Institute of Harvard and MIT, Cambridge, MA 02142, USA; Department of Biological Chemistry & Molecular Pharmacology, Harvard Medical School, Boston, MA, 02115, USA; Department of Biostatistics and Translational Medicine, Medical University of Łódź, Łódź, Poland; Department of Medical Oncology, University Medical Center Groningen, Groningen 9713 GZ, the Netherlands

## Abstract

p53-binding protein 1 (53BP1) is a key mediator of the DNA damage response and genome stability. While its interphase function is well-characterized, its mitotic role remains less understood. Here we show that aberrant activation of 53BP1 through the loss of its negative regulator, TIRR, leads to mitotic abnormalities including altered spindle geometry, kinetochore-microtubule (k-MT) attachment errors and whole chromosome missegregation. We demonstrate that loss of TIRR results in excess interaction between 53BP1 and the key mitotic kinase Polo-like kinase 1 (PLK1), altering PLK1’s activation, spatial distribution, and its interaction with known PLK1 substrates at multiple mitotic stages. Moreover, due to PLK1’s established role in CENP-A loading, hyperactivation of 53BP1 compromises CENP-A loading, triggers gradual loss of CENP-A from centromeres and generates severe kinetochore assembly defects. These findings uncover a non-canonical mitotic function of 53BP1 as a key regulator of PLK1 activity and chromosome segregation fidelity.

## Introduction

Genome stability relies on the coordinated action of pathways that repair DNA lesions during interphase and ensure accurate chromosome segregation during mitosis^1^. Central to this process is the DNA damage response (DDR), an intricate network that detects, signals, and repairs diverse forms of DNA damage. Among DDR factors, p53-binding protein 1 (53BP1) plays a pivotal role in maintaining genome stability by directing double-strand break (DSB) repair pathway choice and modulating p53-dependent transcription^2^. 53BP1 also contributes to class switch recombination^3^ and determines PARP inhibitor sensitivity in BRCA1-deficient tumors ^4, 5^.

53BP1 function is tightly regulated to prevent inappropriate activation outside its canonical role in DNA repair. In interphase, the Tudor-Interacting Repair Regulator (TIRR) binds the 53BP1 tandem Tudor domain to block its interaction with H4K20me2^6^ and dimethylated p53 (K382me2), thereby restraining its repair and transcriptional functions^7^. Structurally, TIRR engages the aromatic cage of the 53BP1 Tudor domain and thereby prevents binding to H4K20me2-marked nucleosomes and K382me2-p53, two key effector interactions that underlie 53BP1’s functions in DNA repair and p53 activation^8^. Thus, TIRR acts as a molecular rheostat that calibrates 53BP1 activity in response to cellular context; its loss is therefore expected to broadly deregulate 53BP1 function beyond the well-characterized interphase setting.

At the onset of mitosis, 53BP1 is phosphorylated at T1609 and S1618, which leads to its dissociation from DNA damage foci^9^, contributing to inactivation of canonical DNA double-strand breakrepair pathways during mitosis. We have previously demonstrated that forced retention of 53BP1 at DNA damage foci during mitosis leads to genomic instability, manifesting as micronuclei and lagging chromosomes^9^. These findings imply that unscheduled activation of 53BP1’s canonical DSB repair function during mitosis is detrimental for chromosome segregation^10^.

Despite the clear inactivation of 53BP1’s DNA repair function during mitosis, accumulating evidence suggests that 53BP1 performs distinct, non-canonical functions in this phase of the cell cycle. 53BP1’s most studied mitotic function is its role in the “mitotic timer” that culls cells that, because of mitotic defects, undergo prolonged mitosis and activate p53-mediated arrest or death in the ensuing interphase^11–13^. This is achieved by the formation of a ternary stopwatch complex, consisting of 53BP1, USP28 and p53, requires PLK1-mediated phosphorylation of 53BP1^14^, and is retained over successive generations^15^.

In addition to its function in mitotic surveillance, 53BP1 has been proposed to play several additional, less well-characterized roles in mitosis. Although 53BP1 does not generally associate with chromatin during mitosis^16^, 53BP1 is reported to localize to the outer kinetochore from prophase through metaphase and play a role in regulating spindle-kinetochore attachment^17^. In addition, it has been proposed to localize to centrosomes and associate with deubiquitase USP7 to regulate centrosome positioning to maintain bipolarity^18^. Moreover, 53BP1 interacts with the anaphase-promoting complex/cyclosome (APC/C) coactivators CDC20 and CDH1, acting as an APC/C inhibitor; in turn, APC/C promotes 53BP1 degradation during anaphase, forming a feedback loop critical for maintaining mitotic homeostasis under stress conditions^19^.

Despite these observations, the mechanistic role of 53BP1 in mitosis remains incompletely understood. We hypothesized that loss of TIRR could aberrantly “activate” 53BP1 in mitosis, revealing a previously unrecognized dimension of its function. Indeed, TIRR knockout cells displayed striking mitotic defects—centric lagging chromosomes, micronuclei, spindle assembly checkpoint activation, altered spindle geometry, and kinetochore assembly errors—all of which were fully rescued by 53BP1 depletion. These findings demonstrate that aberrantly activated 53BP1, rather than loss of TIRR itself, underlies these mitotic abnormalities. Mechanistically, aberrantly activated 53BP1 interacts excessively with PLK1, disrupting PLK1’s localization and phosphorylation status, and its interactions with key mitotic substrates, including those required for CENP-A deposition. Together, these findings uncover a DNA repair–independent mitotic role for 53BP1 in orchestrating PLK1 activity and highlight an unanticipated regulatory axis between TIRR, 53BP1, and PLK1 that safeguards chromosomal stability.

## Results

### Aberrant activation of 53BP1 disrupts spindle-kinetochore attachment, triggering SAC activation

We used TIRR CRISPR/Cas9 knockout (ΔTIRR) or siRNA depletion of TIRR to study 53BP1 activation in mitosis. Loss of TIRR triggered the formation of lagging (ACA/kinetochore-positive) chromosomes (Fig. 1a-d) that result from the missegregation of intact chromosomes during mitosis. As expected for mitotic segregation errors, ΔTIRR and TIRR siRNA cells showed abnormal postmitotic nuclear morphology as evidenced by a decrease in circularity and solidity (Fig. 1e, f). This aberrant activation of 53BP1 in the absence of TIRR did not affect 53BP1’s mitotic surveillance function. In contrast to USP28 siRNA treatment, loss of TIRR did not result in alteration of p53 levels during prolonged mitotic arrest. (Supp fig. 1a).

**Figure 1:**
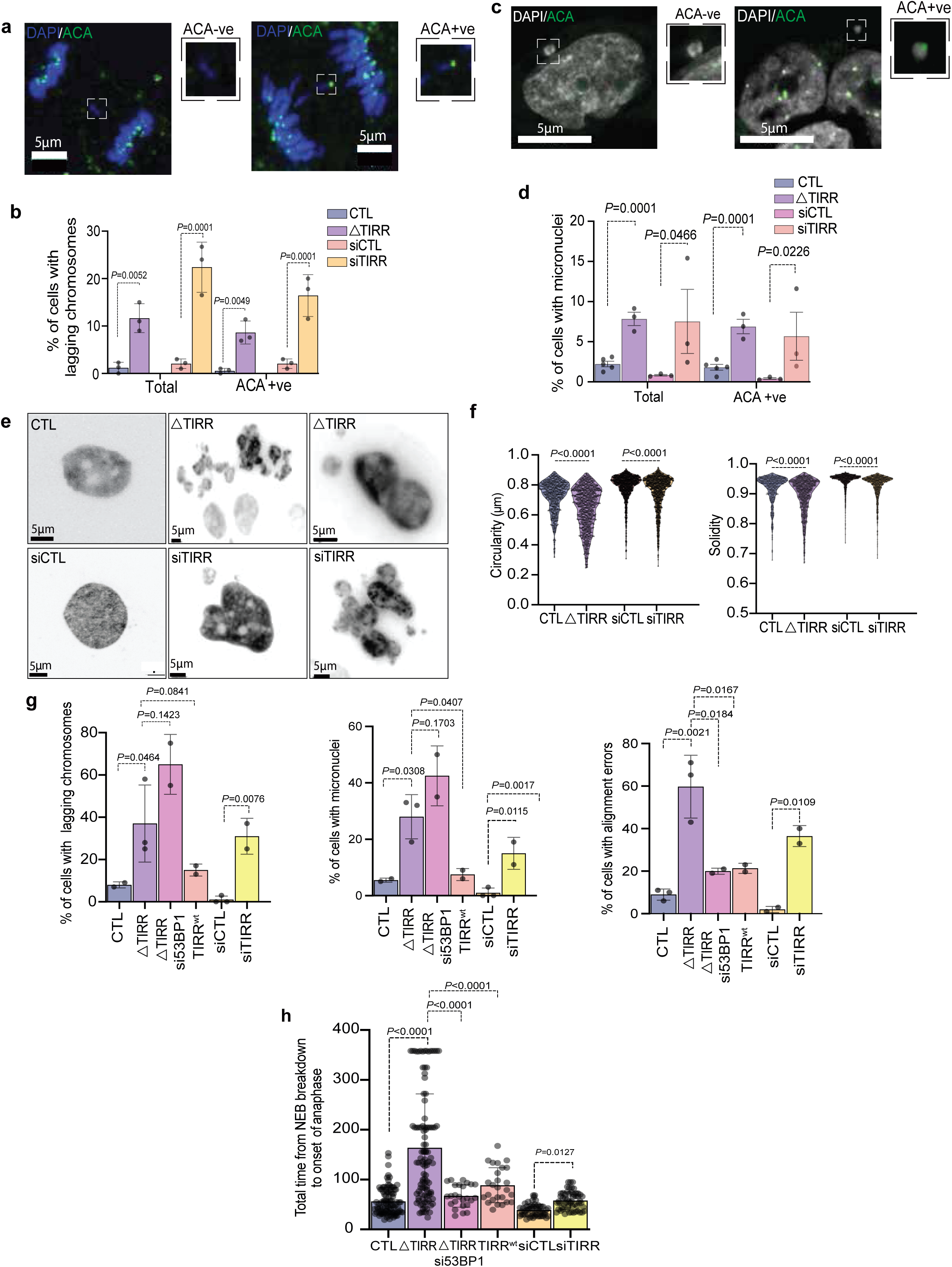
Loss of TIRR leads to mitotic abnormalities. Representative images of RPE-1 cells (control (CTL)), ΔTIRR, siCTL, siTIRR) exhibiting either ACA-positive (centric) or ACA-negative (acentric) (**a**) lagging chromosomes and (**c**) micronuclei (MN), Quantification of the percentage of cells with (**b**) lagging chromosomes and (**d**) MN (centric/acentric). (**e**) Representative images of DAPI staining showing nuclear morphology of RPE-1 control (CTL), ΔTIRR, siCTL and siTIRR cells, fixed 6 hours post-release from RO-3306-induced arrest at G2/M. (**f**) quantification of nuclear circularity and solidity for the experiment in **(e). (g)** Bar graphs showing the frequency of mitotic errors (lagging chromosomes, MN, alignment errors) visualized by high resolution live cell imaging of cells progressing through mitosis. **(h)** Bar graph depicting time from NEBD to anaphase onset determined by live cell imaging of RPE-1 cells as indicated expressing mCherry-H2B and GFP-α-Tubulin, initiated following the release from RO-3306. Data in (**b, d, g**) are mean+/-S.E from three independent experiments (**b** n = 150 anaphase cells and **d** n =1000 cells). Data in (**f**) is represented as violin plot (n =1000 cells) from one representative experiment with three biological replicates. Bar graphs (mean+/-S.D) **(h)** represent mitotic cells visualized in three independent time-lapse imaging experiments (n > 35 cells each). Unpaired, two-tailed student’s t-test results as indicated. Scale bars as depicted.

As a first step to determine the cause of whole chromosome mis-segregation in TIRR-depleted conditions, we used time-lapse microscopy of cells expressing fluorescent histone (mCherry-H2B) and microtubules (GFP-α-Tubulin). TIRR-deficient cells exhibited a significant increase in chromosome alignment errors, resulting in anaphase lagging chromosomes and micronuclei (Fig. 1g, Supp fig. 1b). Because chromosome attachment errors can lead to spindle assembly checkpoint (SAC) activation, we measured the time from nuclear envelope breakdown to anaphase onset. We observed a striking delay in mitotic progression in ΔTIRR and a mild delay in TIRR siRNA cells compared with control cells (Fig. 1h). SAC activation as the cause of the mitotic delay in ΔTIRR cells was confirmed by increased MAD1 signal on kinetochores in metaphase (Supp fig. 1b, c).

These mitotic defects could arise either from an 53BP1-independent function of TIRR or from its regulation of 53BP1. To distinguish between these possibilities, we first examined where TIRR and 53BP1 interact in mitosis using proximity ligation assay (PLA). PLA signal was present exclusively in the nucleocytoplasm from prophase to telophase (Supp fig. 2a, b). This suggests that TIRR sequesters 53BP1 in the nucleoplasm, preventing association with kinetochores during mitosis. These mitotic defects are a direct consequence of 53BP1 aberrant activation rather than an independent function of TIRR, because both overexpression of wild-type TIRR (TIRR^WT^) and depletion of 53BP1 in ΔTIRR RPE-1 cells rescued chromosome alignment defects (Fig. 1g, Supp video file) and mitotic delay (Fig. 1h). As expected, overexpression of TIRR^WT^ or depletion of 53BP1 did not rescue pre-existing nuclear abnormalities (micronuclei and lagging chromosomes), which are irreversible once formed.

### During mitosis, TIRR keeps excess 53BP1 away from the fibrous corona

We next assessed 53BP1 localization at the kinetochore using super-resolution microscopy. 53BP1 in both CTL and ΔTIRR cells co-localized closely with CENP-E, a marker of fibrous corona that assembles on the outer kinetochore at the onset of mitosis and is involved in spindle-kinetochore attachments. In contrast, 53BP1 did not co-localize with the inner kinetochore marker ACA (Supp fig. 3a, b). Additionally, treatment with the MPS1 inhibitor empesertib, which leads to a loss of fibrous corona without affecting the rest of the assembled kinetochore^20–22^, resulted in complete disappearance of 53BP1 signal (Supp fig. 3c), providing further evidence that 53BP1 localizes to the fibrous corona, which confirms previous observations^17^.

Quantitative analysis of 53BP1 kinetochore signal in control and ΔTIRR cells showed substantially higher levels of kinetochore-associated 53BP1 upon TIRR loss (Supp fig. 3c, d). Higher levels of 53BP1 were also observed in both soluble and insoluble fractions of mitotically-synchronized ΔTIRR cells (Fig. 2a). 53BP1 intensity peaked at the kinetochore in prophase to metaphase but significantly decreased at the onset of anaphase as reported previously^18^ (Fig. 2b, c). Consistent with this, there was a higher co-occurrence of 53BP1 with kinetochore marker ACA in ΔTIRR cells compared to control (Supp fig. 3e).

**Figure 2:**
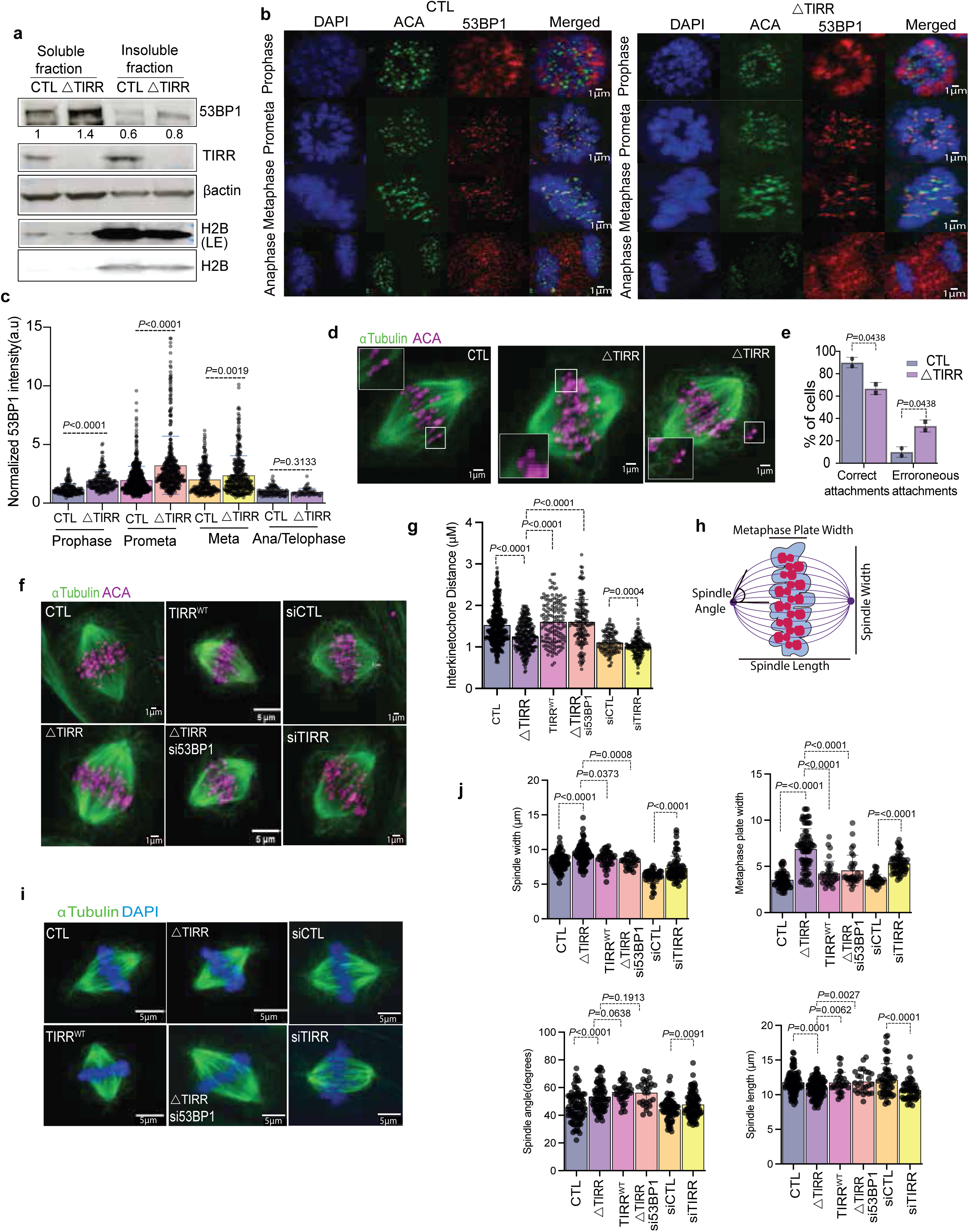
53BP1 dysregulation results in increased 53BP1 localization to kinetochore, altered spindle geometry k-MT attachment errors. **(a)** Immunoblots of cell fractionated lysates from nocodazole-synchronized cells showing protein levels as indicated. **(b)** Immunostaining of RPE-1 cells post-release from RO-3306 with antibodies as indicated, **(c)** normalized 53BP1 intensities of CTL and Δ53BP1 at the mitotic stages as indicated. **(d)** Immunostaining of RPE-1 CTL and Δ53BP1 cells synchronized by RO-3306 release into MG132 for 1 hour with antibodies as indicated. **(e)** Quantitation of correct and erroneous attachments in **d**. **(f)** Representative images of the indicated RPE-1 cells synchronized by the release from RO-3306 into 20µM proTAME for 2 hours and stained with ACA and α-Tubulin antibodies. **(g)** Quantification of inter-kinetochore distance^23^ defined as the distance between two sister kinetochores based on ACA staining (n > 150 kinetochore pairs). **(h)** Schematic depicting measurements of spindle geometry parameters. (**i)** Representative images of the cell types as indicated treated similarly as in **(c)** and stained with α-Tubulin and DAPI**. (j)** Quantification of spindle length, angle, width and metaphase plate width. Data in **c, g, j** is represented as mean+/-S.D (n >150 kinetochores in **c** and **g, j** =n>35 mitotic cells) from at least two independent experiments. Data in (**e**) are mean+/-S.E from two independent experiments (n >150 kinetochores). Unpaired, two-tailed student’s t-test as indicated. Scale bars as depicted.

Because TIRR interaction with 53BP1 blocks its ability to bind H4K20me2 marked chromatin at DNA damage sites^6, 8^, excess 53BP1 at kinetochores might be due to its recruitment to H4K20me2 in the absence of TIRR. However, H4K20 dimethyl marks were present throughout mitotic chromatin (Supp fig. 3e) and were not restricted to the centromere, making it unlikely that 53BP1 recruitment to kinetochores was dependent on H4K20me2. To test the possibility that 53BP1 is recruited to sites of centromeric DNA damage, we irradiated mitotically-synchronized RPE-1 cells and performed colocalization analysis with γ-H2AX and centromeric sequences marked by ACA labeling. There was minimal overlap between 53BP1 and γ-H2AX foci in both control and in ΔTIRR cells. By contrast, the overlap between 53BP1 and centromeres was unaffected by irradiation. (Supp fig. 3f, g, h).

Together, these data show that loss of TIRR causes excess accumulation of 53BP1 at the outer kinetochore from prophase to metaphase in a manner independent of DNA damage, supporting the conclusion that 53BP1 performs a mitotic function distinct from its role in DNA repair.

### Loss of 53BP1 control leads to altered biophysical properties of mitotic spindle

We next investigated the cause of higher chromosome mis-segregation rates upon the loss of TIRR. We found a higher frequency of cells with kinetochore-microtubule (k-MT) attachment errors in ΔTIRR cells compared to controls (Fig. 2d, e). Erroneous k-MT attachments may also affect MT tension forces, which can be inferred from the inter-kinetochore distance^23^. Indeed, we found that ΔTIRR and siTIRR showed decreased IKD compared to CTL cells (Fig. 2f, g), providing further evidence that aberrant activation of 53BP1 leads to k-MT errors. To characterize spindle geometry defects, we performed live cell imaging of cells released into mitosis in the presence of APC/C inhibitor proTAME, which arrests the cells in metaphase. The mitotic spindle was quantified as illustrated in Fig. 2h. Quantitative analysis showed that loss of TIRR led to a significant increase in the spindle and metaphase plate width, and a significant decrease in spindle angle and length (Fig. 2i, j). These findings demonstrate that 53BP1 regulation by TIRR is essential to maintain proper spindle architecture and facilitate proper k-MT attachment. Importantly, both IKD and spindle geometry defects were rescued by 53BP1 depletion or re-expression of wild-type TIRR in ΔTIRR cells, supporting our conclusion that these defects are due to 53BP1 dysregulation rather than a 53BP1-independent function of TIRR.

### 53BP1 hyperactivation leads to kinetochore assembly defects by blocking CENP-A loading

Spindle-kinetochore attachment errors could arise as a consequence of kinetochore assembly defects. To examine this possibility, we assessed the levels of both inner and outer kinetochore proteins upon the loss of TIRR. Strikingly, loss of TIRR resulted in a significant reduction in the levels of both inner kinetochore (CENP-A, CENP-C, CENP-T) and outer kinetochore (HEC1) proteins (Fig. 3a, b).

**Figure 3:**
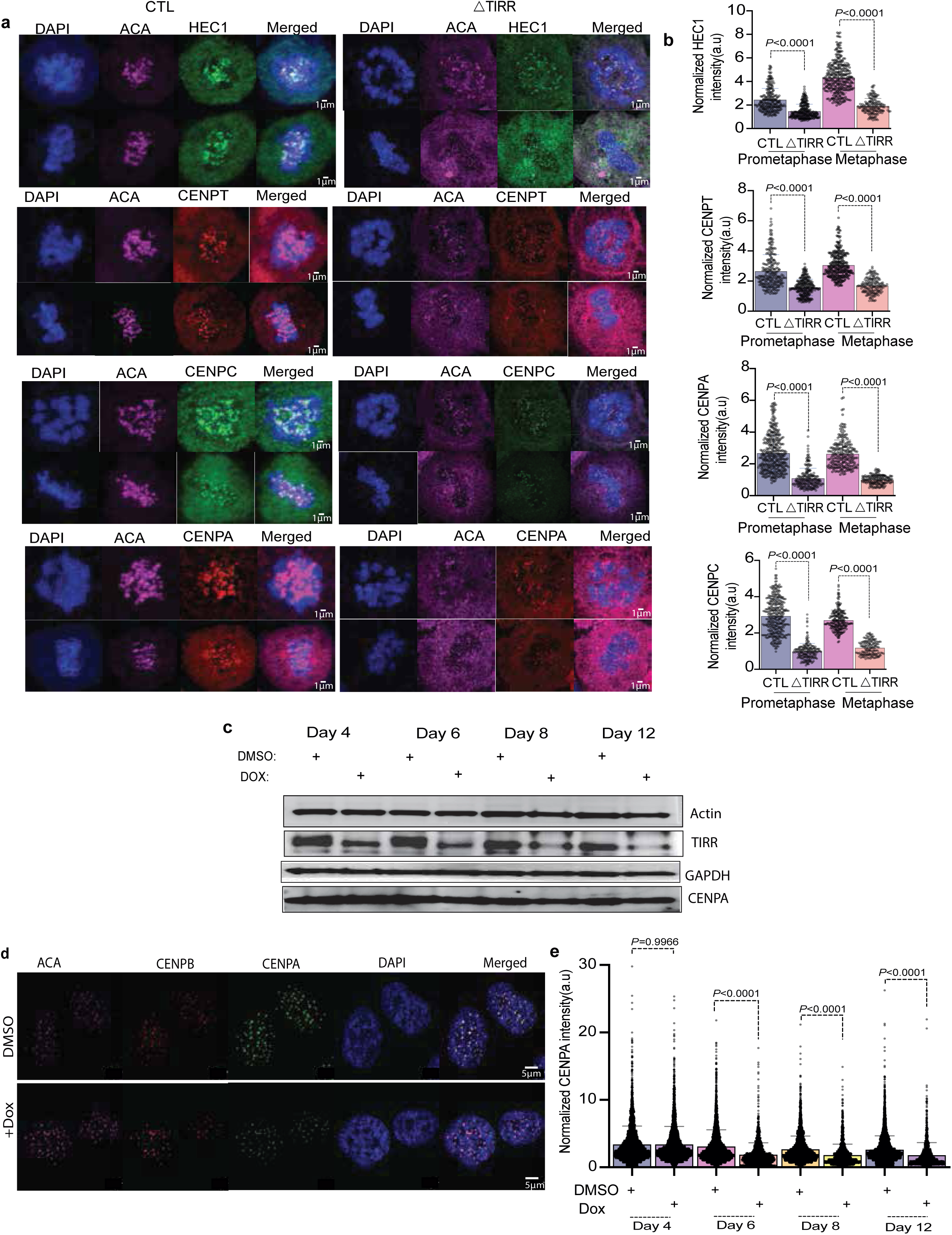
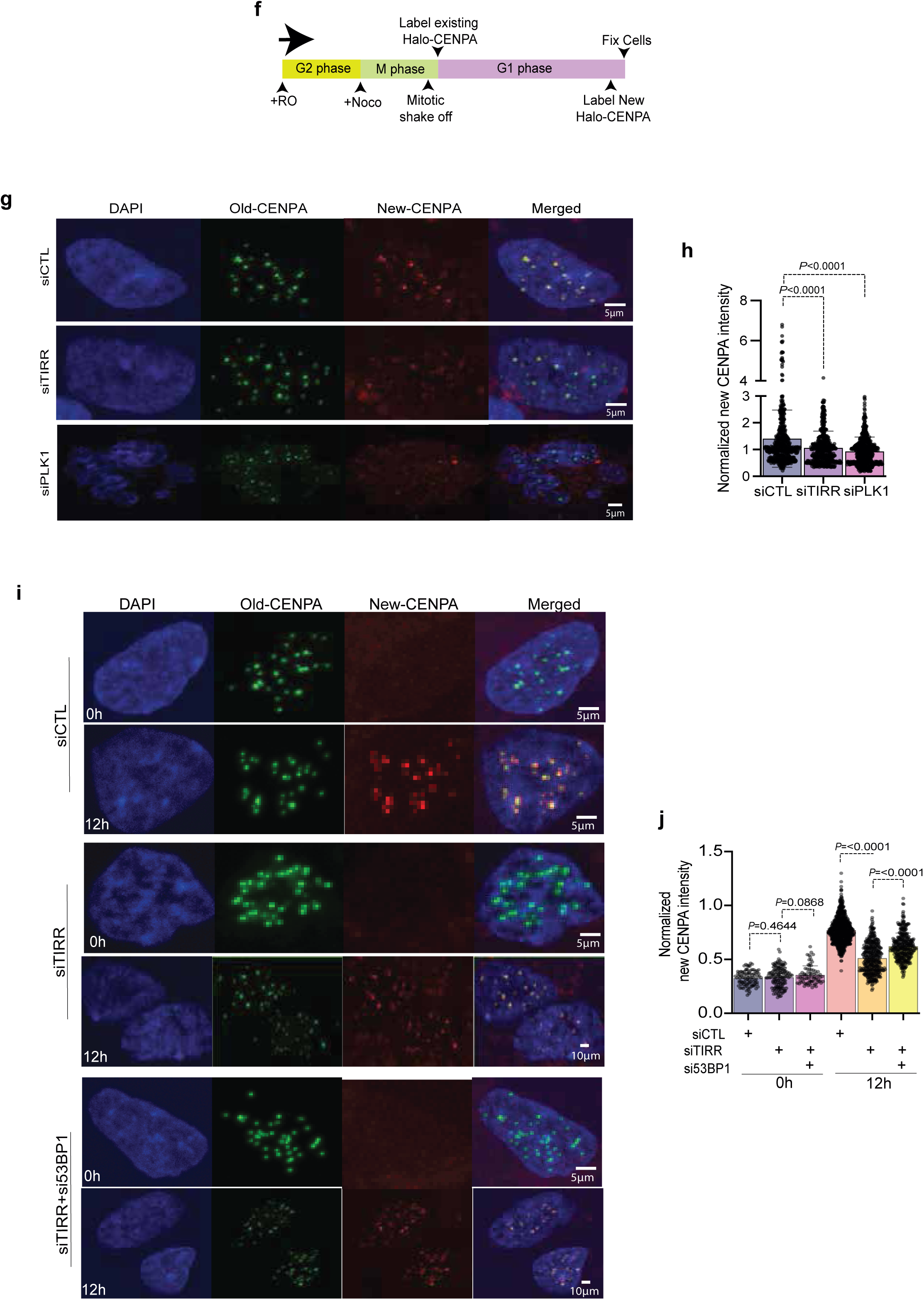
Loss of TIRR impairs CENP-A loading and leads to kinetochore assembly defects. **(a)** Immunostaining of RPE-1 CTL and ΔTIRR cells with the indicated antibodies in prometaphase (*top row*) and metaphase (*bottom row*). **(b)** Normalized intensities of the indicated proteins at kinetochores. **(c)** Immunoblots showing levels of proteins as indicated upon DMSO or doxycycline treatment in inducible shTIRR RPE-1 cells. **(d)** Representative images of doxycycline-inducible shTIRR RPE-1 cell fixed on day 6 and stained with the indicated antibodies. **(e)** Quantification of normalized CENP-A fluorescence intensity on the indicated day after initiation of doxycycline treatment. **(f)** Schematic depicting Halo-tag CENP-A incorporation assay in RPE-1 cells. **(g, i)** Representative images of pre-existing (green) and newly loaded CENP-A in cells treated as indicated. **(h, j)** Quantification of newly incorporated Halo-Tag-CENP-A fluorescence intensity in cells as indicated. Data in (**b-j**) is represented as mean+/-S.D (n >200 kinetochores) and experiment was repeated two times. Unpaired, two-tailed student’s t-test as indicated. Scale bars as depicted.

Because 53BP1 localizes to the outer kinetochore, it is unlikely to disrupt localization of inner kinetochore proteins directly. This raised the possibility that 53BP1 hyperactivation may instead interfere with kinetochore assembly over successive cell cycles. To test this possibility, we introduced doxycycline-inducible short hairpin (shTIRR) in RPE-1 cells and confirmed inducible depletion of TIRR (Fig. 3c, Supp fig. 4a).This enabled temporal control of TIRR depletion and assessment of kinetochore assembly defects over multiple cell divisions. We found that CENP-A intensity at centromeres was not significantly reduced in the first 4 days following doxycycline induction, but reduction became apparent and significant from day 6 onward (Fig. 3c-e). CENP-B levels showed the reverse pattern, with a significant increase by day 6 consistent with prior reports^24–26^(Supp fig. 4c). This gradual loss of CENP-A over multiple cell cycles was suggestive of a CENP-A deposition defect rather than acute loss over a single cell cycle.

To maintain constant CENP-A levels at kinetochores, centromeric CENP-A levels are replenished in G1 phase of every cell cycle by a well-orchestrated deposition process ^27, 28^. To test whether kinetochore assembly defects in TIRR-depleted cells occurred as a result of the perturbed CENP-A deposition, we employed an established CENP-A deposition assay by sequential labeling of pre-existing and newly incorporated CENP-A in RPE-1 cells with Halo-tagged CENP-A at its endogenous locus^29^. Pre-existing CENP-A was labeled in nocodazole-arrested mitotic cells. Cells were then released and allowed to progress into G1 in the presence of a second dye to label newly incorporated CENP-A (Fig. 3f). We observed a significant reduction in newly incorporated CENP-A after 12 hours in TIRR-depleted cells compared to controls (Fig. 3g, h). The degree of reduction was comparable to PLK1 siRNA, which is known to abrogate CENP-A deposition^27, 30, 31^. Together, these results indicated that dysregulation of 53BP1 leads to impaired CENP-A deposition, causing progressive reduction in CENP-A levels that ultimately trigger kinetochore assembly defects once CENP-A levels fall below a certain threshold. Finally, we tested whether co-depletion of 53BP1 could rescue the CENP-A deposition defect in TIRR kd cells, using endogenously tagged Halo-CENP-A cells as described earlier. Indeed, we observed significant rescue of CENP-A deposition upon 53BP1 loss (Fig. 3i, j), indicating that the CENP-A deposition defect is attributable to aberrant 53BP1 activation.

### 53BP1 interacts with PLK1 and regulates its localization and function in mitosis

To identify the mechanism responsible for CENP-A deposition defect, we immunoprecipitated 53BP1 from mitotically-synchronized control and ΔTIRR lysates and identified interacting proteins by mass spectrometry (Supp fig. 5 a, b). Intriguingly, we found increased association of 53BP1 with PLK1, a key mitotic kinase essential for multiple stages of cell division, including spindle formation, chromosome attachment, and cytokinesis^32–34^. We validated increased association between 53BP1 and PLK1 in the absence of TIRR using PLA (Fig. 4a) and IP-immunoblot (Fig. 4b). Interestingly, the 53BP1-PLK1 interaction was enriched throughout the nucleocytoplasm, rather than on chromatin. We next tested whether increased 53BP1-PLK1 association alters PLK1 interaction with its known substrates. Using IP–immunoblot, we observed increased interaction between PLK1 and its known substrates CENP-F and BUBR1 in ΔTIRR cells (Fig. 4c, d). Importantly, this enhanced association was rescued by 53BP1 depletion (Fig. 4d), confirming a direct effect of excess 53BP1-PLK1 interaction.

**Figure 4:**
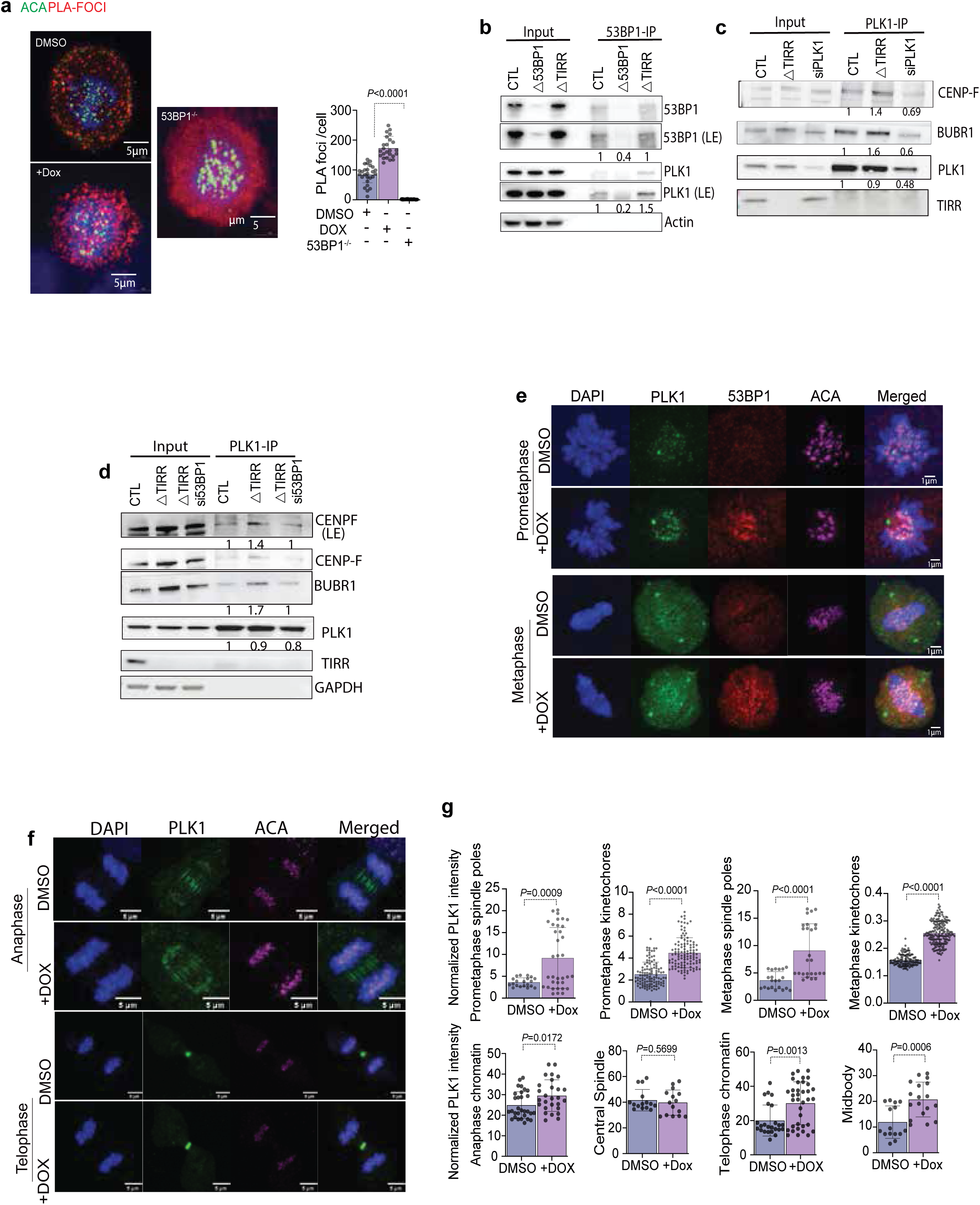
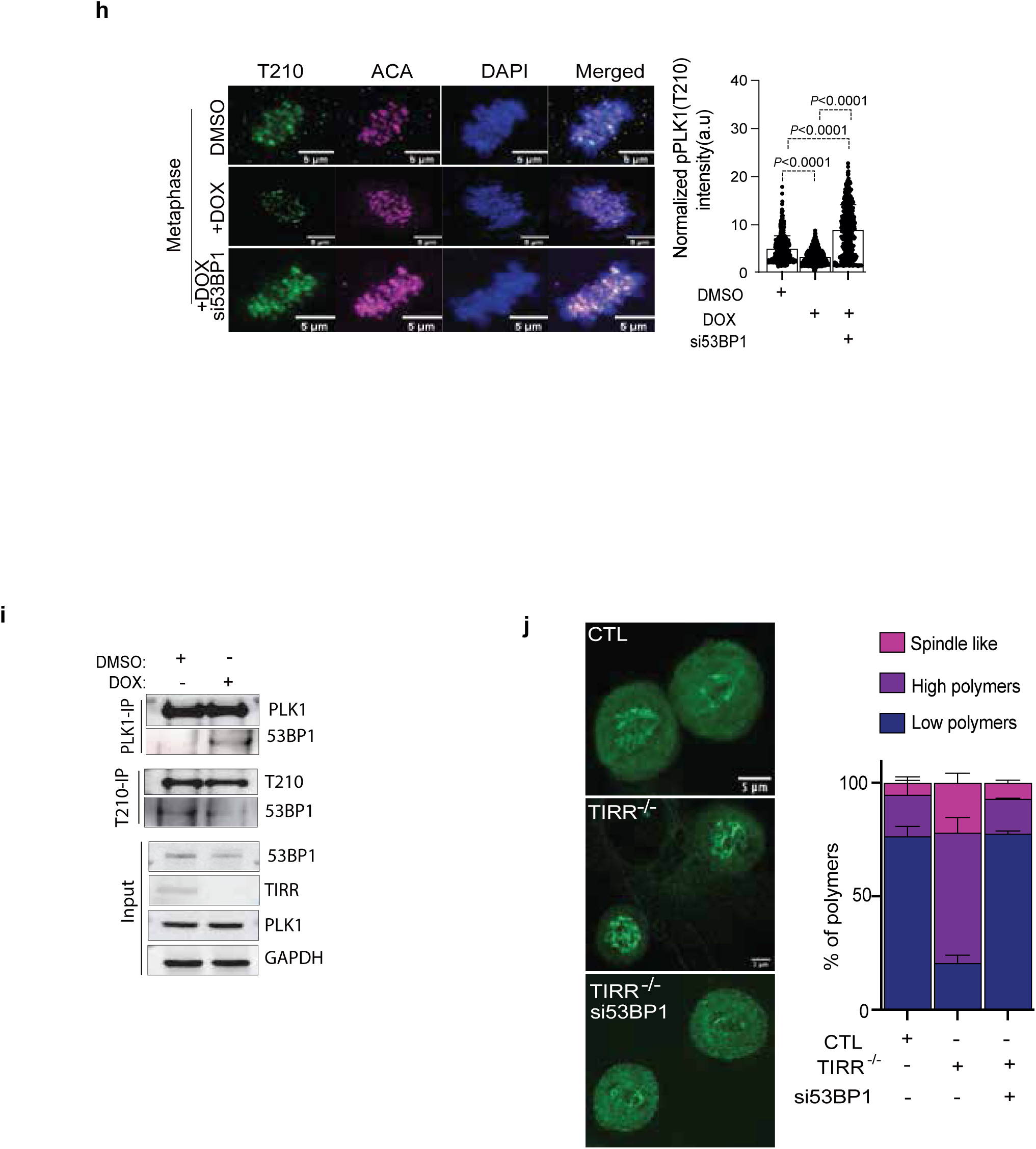
53BP1 regulates PLK1 localization and function. **(a)** Representative images of proximity ligation assay between 53BP1 and PLK1 performed in control (DMSO), TIRR shRNA treated with doxycycline and Δ53BP1 synchronized in prometaphase by nocodazole. Graph (*bottom right)* depicts the number of PLA foci observed in cells as indicated (CTL, TIRR shRNA treated with doxycycline and Δ53BP1. **(b)** 53BP1 immunoprecipitation from mitotically-synchronized cell lysates as shown immunoblotted with the indicated antibodies. **(c, d)** Immunoblots showing BUBR1, PLK1 and CENP-F interaction after PLK1 immunoprecipitation from nocodazole-synchronized lysates as shown. **(e, f, h)** Representative images of prometaphase and metaphase RPE-1 cells treated as indicated and stained with the antibodies as shown. Normalized intensities of **(e, f)** total PLK1 and **(h)** phospho-PLK1(T210). **(i)** Immunoblots of PLK1 and phospho-PLK1(T210) immunoprecipitation from mitotically-synchronized lysates blotted with the antibodies as shown. **(j)** Representative images of metaphase RPE-1 cells depicting indicated spindle microtubule polymer levels, as described in a previous study^62^, after the cells had been exposed to cold shock treatment; quantification of the frequency of indicated spindle microtubule polymer levels observed after cold shock treatment (right). Data in (**a-j**) is represented as mean+/-S.D (n>35 mitotic cells; n >200 kinetochores) and experiment was repeated at least two times. Unpaired, two-tailed student’s t-test as indicated. Scale bars as depicted.

PLK1 is spatially and temporally regulated in distinct subcellular localization patterns throughout mitosis^33–38^. Although numerous PLK1 substrates have been identified^39, 40^, the mechanisms governing its spatiotemporal localization and precise functional regulation remain only partially understood. To determine whether enhanced interaction with 53BP1 alters its cellular localization, we analyzed PLK1 localization across different mitotic stages. Notably, we observed significant alterations in PLK1 localization across various mitotic subcellular compartments upon TIRR depletion (Fig. 4e, f). PLK1’s accumulation at kinetochores and spindle poles was markedly enhanced, and its presence at chromatin and the midbody was also aberrantly elevated (Fig. 4g). These findings suggest that TIRR depletion alters spatiotemporal localization of PLK1, potentially impairing its role in regulating k-MT attachments and other key mitotic functions.

PLK1 is regulated by phosphorylation on its T210 residue in its T-loop segment, which has been reported to play a key role in mitotic progression and kinetochore function^41–44^. Interestingly, we observed a reduction in T210 phosphorylation at kinetochores in metaphase-synchronized TIRR-depleted cells, which was rescued by 53BP1 co-depletion (Fig. 4h). Moreover, while hyperactivation of 53BP1 results in excessive interaction with PLK1, interaction between 53BP1 and T210 phosphorylated form of PLK1 is diminished (Fig. 4i). This suggests that excessive interaction with 53BP1 leads to localized inhibition of PLK1 on kinetochores, which can compromise its key mitotic functions including spindle-kinetochore attachment regulation as previously reported^45–48^. Because active PLK1 at kinetochores plays an essential role in regulating spindle microtubule dynamics^49, 50^, we tested whether a reduction in the T210 phosphorylated form of PLK1 at kinetochores would alter spindle microtubule stability. Indeed, we observed higher spindle MT stability in TIRR-depleted cells in cold shock assays, which was rescued by 53BP1 co-depletion (Fig. 4j).

### Aberrant activation of 53BP1 impairs PLK1-dependent CENP-A deposition

PLK1 plays an essential role in regulating CENP-A deposition by phosphorylating components of the Mis18 complex^27, 31, 30^. We first tested whether excessive 53BP1-PLK1 interaction in TIRR-depleted cells persisted into G1. Indeed, we observed increased 53BP1-PLK1 PLA foci upon TIRR kd in G1 cells following release from nocodazole for 2 and 4 hours (Fig. 5a, b). We next tested whether TIRR depletion altered PLK1 recruitment to centromeres at this stage of the cell cycle. Indeed, we observed a significant reduction in total and phospho-T210 PLK1 in early G1 upon TIRR kd (Fig. 5c, d).

**Figure 5:**
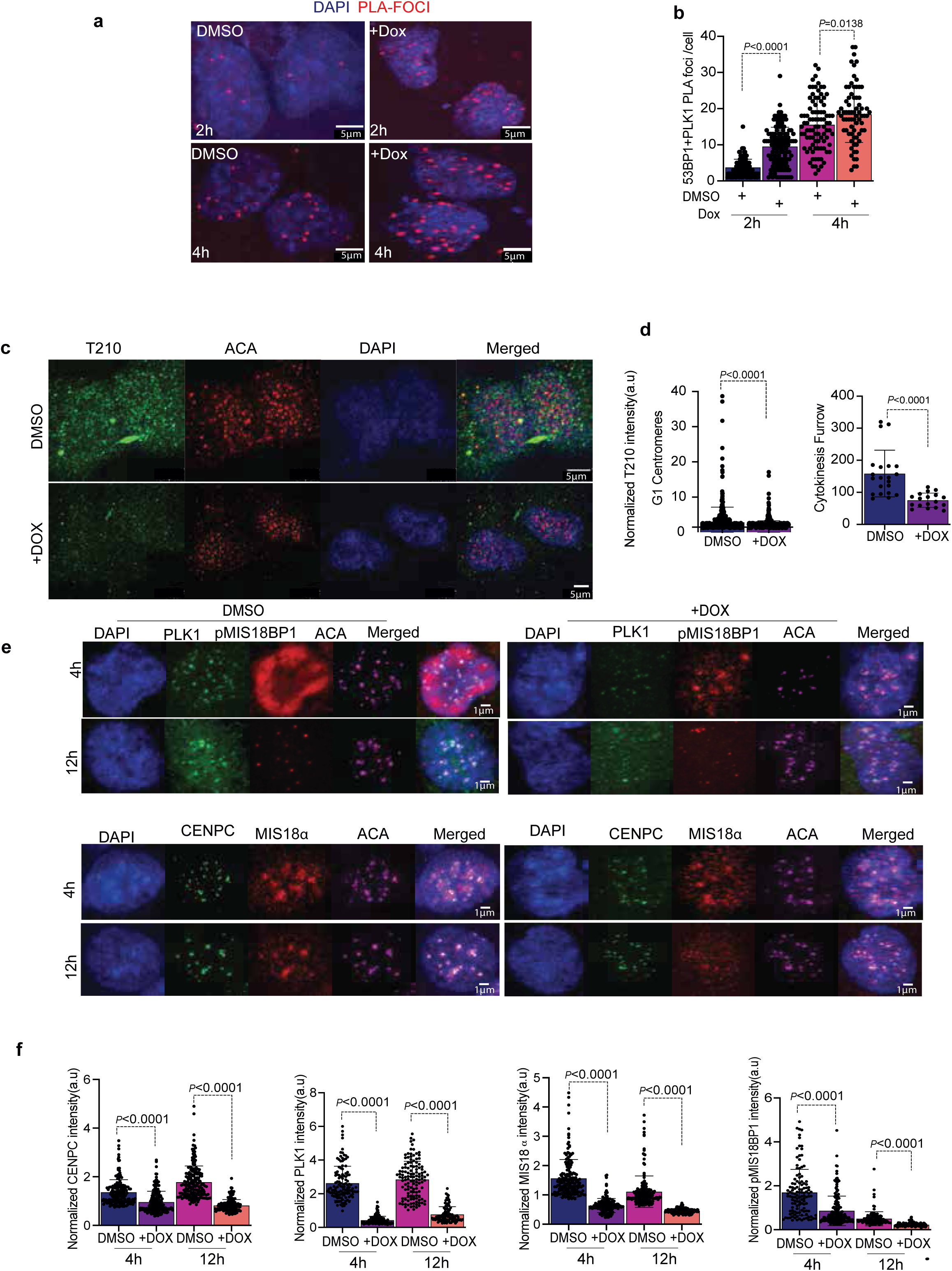
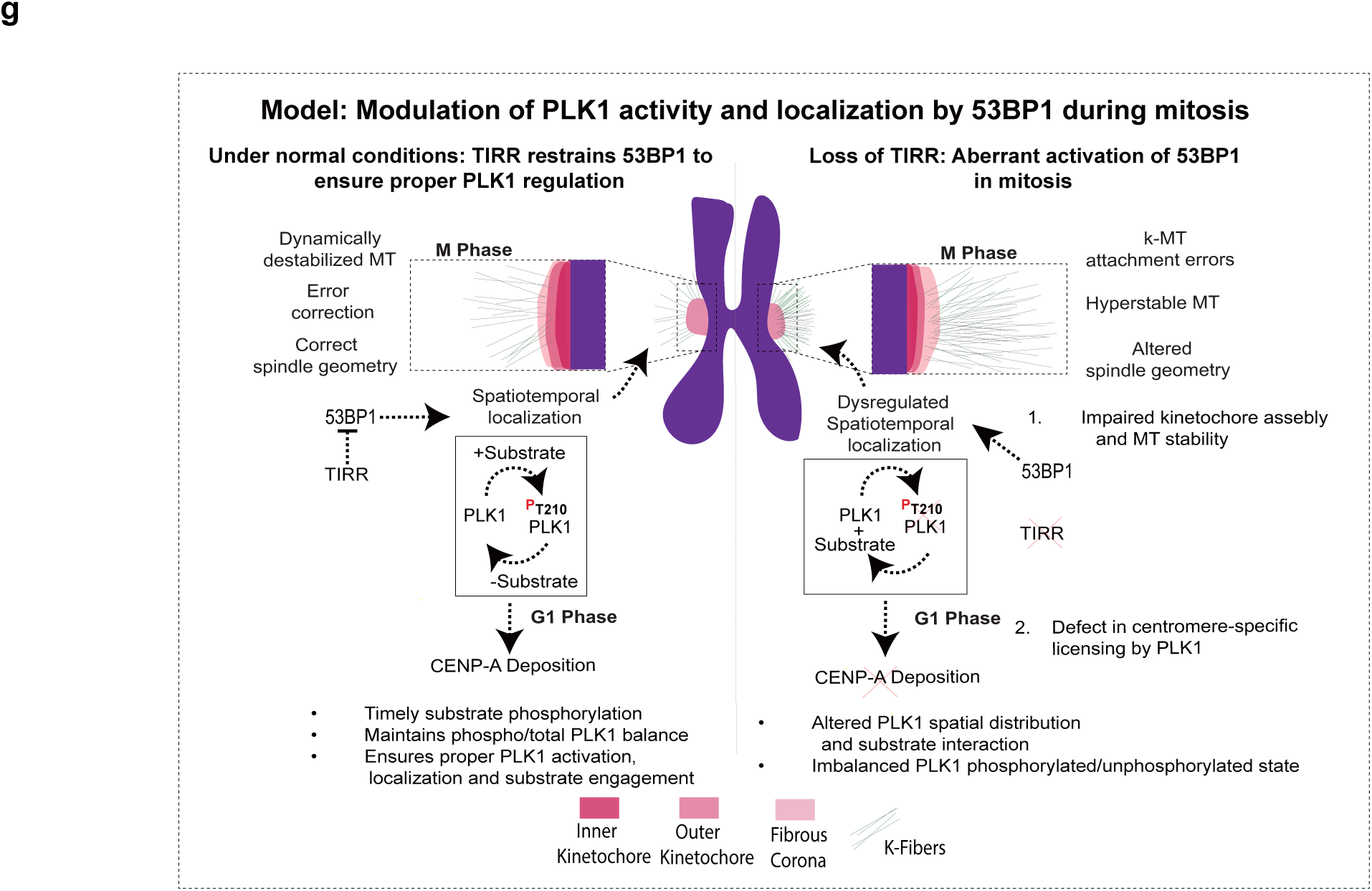
Dysregulated 53BP1 impairs CENP-A deposition. **(a)** Representative images of proximity ligation assay between 53BP1 and PLK1 performed in RPE-1 cells as indicated synchronized by mitotic shake-off from nocodazole and fixed at 2h and 4h post-release. **(b)** Graph depicting the number of PLA foci observed after treatments as indicated. **(c)** Representative images of RPE-1 cells fixed post-release from RO-3306 and stained with the indicated antibodies. **(d)** Normalized intensities of PLK1(T210) at the indicated subcellular locations after treatments as shown. **(e)** Representative images of RPE-1 synchronized by mitotic shake-off into fresh media and fixed at time points as indicated (4h and 18h), stained with antibodies against PLK1, CENPC, MIS18, phospho-MIS18BP1 and ACA. **(f)** Normalized intensities of CENPC, PLK1, MIS18α, and phospho-MIS18BP1 in cells as indicated. (**g**) Proposed model. Data in (**b, d, f**) is represented as mean+/-S.D (**a**=n>500 centromeres; **d**=n >200 kinetochores and n>35 mitotic cells; **f**>150 centromeres) from at least 2 independent experiments. Unpaired, two-tailed student’s t-test as indicated. Scale bars as depicted.

Recent studies elucidated the molecular mechanism underpinning CENP-A deposition^31, 51, 52, 27, 30^. PLK1 regulates deposition by phosphorylating components of the heterotrimeric Mis18 complex consisting of MIS18α, MIS18β and MIS18BP1 proteins, which, in turn, leads to its recruitment to early G1 centromeres facilitating deposition of HJURP-CENP-A^53–60, 27, 30^. To test whether aberrant activation of 53BP1 impairs PLK1-dependent CENP-A deposition pathway, we assessed recruitment of MIS18α/β and phosphorylation of MIS18BP1 on T702 (essential for CENP-A deposition) on centromeres in early G1^27^.

We observed a striking reduction in phospho-T702 MIS18BP1 and MIS18α levels on G1 centromeres after TIRR kd compared to control cells (Fig. 5f). This provides further evidence that hyperactivation of 53BP1 alters PLK1 localization and function and leads to multiple mitotic defects, including impaired CENP-A deposition. In turn, this leads to further exacerbation of mitotic defects over time due to kinetochore assembly failure.

## Discussion

Here, we demonstrate that aberrantly activated 53BP1, a pivotal regulator of genome stability through its role in DNA repair pathway choice and control of p53 levels, results in chromosome segregation errors by excessively binding to PLK1 in the nucleocytoplasm and dysregulating PLK1 activation, localization, and activity during mitosis. This function is independent of its canonical role in DNA repair during interphase and 53BP1’s role in mitotic surveillance and regulation of p53 stability. This dysregulation disrupts spindle geometry and increases the frequency of chromosome attachment errors in the short term. Excessive PLK1-53BP1 binding also impairs CENP-A loading, leading to gradual decline in CENP-A levels over successive cell cycles, ultimately causing severe defects in kinetochore assembly and persistent activation of the spindle assembly checkpoint, which culminates in prolonged mitotic arrest (Fig. 5g).

The clinical significance of these findings is underscored by the frequent dysregulation of 53BP1 and its regulators in human cancer. 53BP1 expression is reduced or lost in a significant fraction of triple-negative breast cancers and other tumor types, and such loss correlates with BRCA1-independent genomic instability^4, 61^. Similarly, TIRR levels are markedly reduced in multiple cancer datasets, suggesting that impaired TIRR-mediated restraint of 53BP1 may contribute to the chromosomal instability (CIN) phenotype observed in these tumors. Our findings raise the possibility that aberrant 53BP1–PLK1 interactions, arising from diminished TIRR expression, impair centromere maintenance and kinetochore assembly and thereby drive CIN in cancer cells. This TIRR–53BP1–PLK1 axis therefore represents a candidate vulnerability in CIN-high tumors, warranting future investigation as a therapeutic target or biomarker of mitotic fidelity.

In this study, we define a mitotic function of 53BP1 revealed under conditions of its aberrant activation following loss of the negative regulator TIRR. Equally important will be to investigate the consequences of 53BP1 depletion during mitosis. Such an approach will require a system that enables acute degradation or inactivation of endogenous 53BP1 specifically at mitotic entry, thereby allowing its mitotic role to be distinguished from its well-established functions in interphase.

PLK1 function depends on tightly controlled spatiotemporal regulation, yet how its localization is dynamically governed—particularly with respect to the interplay between total and phosphorylated PLK1—remains incompletely understood. Here, we define a novel regulatory axis that coordinates PLK1 localization in space and time. We further show that perturbation of TIRR levels disrupts the coupling between T210 phosphorylation and total PLK1 distribution, leading to aberrant, persistent retention of PLK1 at kinetochores and its mislocalization to centromeres. Future studies will also be needed to elucidate how 53BP1 regulates PLK1 post-translational modifications, including phosphorylation at T210, which is critical for several of its mitotic functions. Our findings indicate that 53BP1 preferentially associates with the unphosphorylated form of PLK1 at T210, and that loss of TIRR-mediated regulation reduces T210 phosphorylation levels at the kinetochore, altering spindle microtubule dynamics and compromising spindle-kinetochore attachment fidelity. Future investigations will be required to delineate the underlying mechanisms of this regulatory process.

We and others have previously shown that 53BP1 must disengage from DNA damage sites at the onset of mitosis to safeguard genome stability, a process thought to prevent aberrant DNA repair during cell division. Our finding that 53BP1 also regulates mitotic PLK1 function raises the alternative possibility that its release from DNA damage sites is required to fulfill this essential mitotic role. Future studies will be needed to disentangle these mechanisms and to define the specific contributions of 53BP1 to mitosis under both physiological and pathological conditions.

## Supplementary figure legends

**Supp fig 1: Loss of TIRR leads to increased mitotic errors. (a)** Immunoblots showing levels of proteins upon PLK4 inhibitor centrinone treatment of indicated RPE-1 cells for 48h. **(b)** Time-lapse fluorescence microscopy of RPE-1 cells as indicated expressing mCherry-H2B and GFP-α-tubulin. Punctate square marks mitotic errors such as lagging chromosomes and chromosome bridges. **(c)** Representative images of RPE1 cells fixed at indicated stages post-release from RO-3306 and stained with ACA and MAD1. **(d)** quantification of MAD1 signal detected at kinetochore at prometaphase and metaphase stages. Data in **(d)** is represented as mean+/-S.D (n>150 kinetochores) with 3 biological repeats. Unpaired, two-tailed student’s t-test as indicated. Scale bars as depicted.

**Supp fig 2: TIRR binds to 53BP1 during mitosis. (a)** Representative images of proximity ligation assay between 53BP1 and TIRR performed in RPE-1, post-release from RO-3306 and fixation at different mitotic phases. **(b)** Number of PLA foci observed in different stages of mitosis. Data in **(b)** is represented as mean+/-S.D (n>25 mitotic cells) with two biological repeats. Unpaired, two-tailed student’s t-test as indicated. Scale bars as depicted.

**Supp fig 3: Increased 53BP1 localization to the fibrous corona in the absence of TIRR is independent of DNA damage signaling. (a)** Immunostaining of RPE-1 CTL and ΔTIRR metaphase spreads with the indicated antibodies. **(b)** Line plots showing normalized kinetochore intensity of 53BP1, ACA (inner kinetochore marker), and CENP-F (outer kinetochore marker). **(c)** Representative images of metaphase spreads in the presence or absence of MPS1 inhibitor of RPE-1 cells stained with antibodies against 53BP1, CENP-E and ACA. **(d)** Representative mages of metaphase spreads of RPE-1 control (CTL) and ΔTIRR cells stained with the antibodies against 53BP1 and ACA. (e) *(Left panel*) Normalized intensities of 53BP1 and kinetochore marker ACA, (*Right panel)* Mander’s overlap coefficient between 53BP1 and ACA. **(f)** Representative images of RPE1 cells released from RO-3306, fixed at different mitotic phases and stained with H4K20me2 and ACA antibodies. **(g)** Representative images of metaphase spreads of RPE-1 control (CTL) and ΔTIRR cells after irradiation (0 Gy or 2 Gy as indicated) stained with the indicated antibodies. **(g)** Normalized intensity of 53BP1 at kinetochore (defined by ACA marker) in cells as indicated. **(i)** Mander’s overlap coefficient between (*Top panel*) 53BP1 and ACA, (Below panel) 53BP1 and γH2AX. Data in (**f, d, g, h**) is represented as mean+/-S.D (n >150 kinetochores and/or n> 35 mitotic cells). These are representative of at least two independent experiments.

**Supp fig 4: CENP-A intensity at centromeres gradually decreases and CENP-B intensity increases upon long-term TIRR knock down. (a)** Immunoblots showing levels of proteins upon DMSO or doxycycline treatment of doxycycline-inducible RPE-1 TIRR shRNA cells. **(b)** Representative images of doxycycline-inducible shTIRR RPE-1 cell fixed on day 2 and stained with the indicated antibodies. **(c)** Quantification of normalized CENPB fluorescent intensity. Data in (**c**) is represented as mean+/-S.D (n >200 kinetochores) with two biological repeats. Unpaired, two-tailed student’s t-test as indicated. Scale bars as depicted.

**Supp fig 5: 53BP1 regulates** PLK1 mitotic localization. Volcano plot of mass spectrometry results following immunoprecipitation of 53BP1 from **(a)** CTL and **(b)** ΔTIRR RPE-1 cells synchronized by nocodazole and mitotic shake-off.

## Methods

### Cell culture

All cells were cultured at 37°C in 5% CO2 atmosphere with 100% humidity. HeLa and 293T were cultured in DMEM-high glucose, pyruvate (Invitrogen, 11995-065) supplemented with 10% (v/v) fetal bovine serum (FBS, Invitrogen, 10437028) and 1% Pen-Strep (Invitrogen, 15140-122). Telomerase-immortalized (hTERT) RPE-1 cells were cultured in DMEM-F12 medium (Invitrogen, 21041-065) containing 10% FBS and 1% Pen-Strep.

#### RPE-1 expressing mCherry-H2B and GFP-α-tubulin for live-cell imaging

RPE-1 stably expressing mCherry-H2B and GFP-α-tubulin were generated by lentiviral transduction. Lentiviral production was performed using 293T cells, which were co-transfected with pMD2.G (Addgene #12259), psPAX2 (Addgene, #12260), and lentiviral plasmid pLenti6-H2B-mCherry (Addgene #89766) or lentiviral plasmid pL304-eGFP-α-tubulin (Addgene #64060) using Lipofectamine 3000, as described above. Viral particles harboring mCherry-H2B or GFP-α-tubulin were filtered as described^63^ and applied in 1:1 ratio to RPE-1. Transduced cells were then sorted to isolate population expressing mCherry-H2B and GFP-tubulin gated for 30 to 70% intensity, using fluorochrome 530/30 for GFP and 610/20 for mCherry into fresh DMEM-F12 medium containing 20% FBS.

### Cell cycle synchronization

For synchronization to G2/M, cells were treated with RO-3306 at the final concentration of 9µM for 16-18 hours. Cells were released from RO-3306-mediated G2/M arrest by wash with pre-warmed 1X phosphate buffer saline (PBS; Corning 21-040) five times before adding fresh media. For synchronization in prometaphase, cells were treated with nocodazole (Sigma Aldrich, M1404) at the final concentration of 100 ng/ml for 6 hours. Cells were released from nocodazole-mediated prometaphase arrest by mitotic shake-off and 4 washes with pre-warmed fresh media. For synchronization in metaphase, cells were at first synchronized in G2/M by treatment with RO-3306. cells were then washed and released into media containing 20µM of proTAME (Medchem Express HY-124955) for 1 hour.

### Image and nuclear shape analysis

For fixed cells, images were acquired with 20x objective as mentioned^63^. Quantitative analysis of images was performed using ImageJ/Fiji. Region of interest (ROI) was either manually drawn around cell boundaries (for mitotic cells) or defined using image segmentation (for nuclei). For nuclear segmentation, either Li or Otsu thresholding was employed on maximum intensity projections of Z-stacks of images acquired using 405nm laser (DAPI-stained). After the images were converted into binary, the objects (nuclei) were processed using “Erode” and “Dilate” processing tools in ImageJ/Fiji. Lastly, the “Watershed” tool was applied to separate the clumped nuclei. The ROIs generated were then overlayed on the 405nm images and used to measure circularity and solidity using Image J/Fiji shape descriptor tools. For measuring nuclear parameters, cells were imaged with a 20x objective in 8×8 tiles. The tiles were stitched and at least 200 cells were analyzed per image. Lobular, fragmented nuclei or nuclei containing blebs, folds and/or crevices were manually scored as abnormal. For quantification of fluorescence intensity of labeled proteins, ROIs were defined as described above and either total (mean intensity * total area) or normalized intensity was determined in the given channel following background subtraction (defined by ROI drawn outside the cell). Normalization was performed to total DAPI intensity. The Manders’ overlap coefficients were derived using the JACoP Fiji tool for colocalization analysis. The thresholding was kept constant throughout the images analyzed.

### Immunofluorescence microscopy

Cells released from synchronization as described above were fixed with 4% paraformaldehyde (Electron Microscopy Sciences, 15710) for 15 minutes. Fixed samples were washed with PBS, permeabilized for 5-10 minutes at room temperature with PBS with 0.5% Triton X-100 and washed with PBS. Samples were then blocked in PBS with 3% BSA (Sigma, 10735086001) for one hour, incubated with primary antibodies diluted in blocking buffer overnight. The samples were washed three times with PBS with 0.05% Triton X-100, incubated with secondary antibodies diluted in blocking buffer at room temperature for 1 hour, and washed again three times in PBS with 0.05% Triton X-100. DNA was stained using Hoechst 33342 for 20 mins, washed and mounted in ProLong Gold antifade (Life Technologies) on glass slides. Alternatively, post-secondary antibody incubation and washes, samples were washed once in Ultrapure Distilled Water (Invitrogen, 10977) before invert-mounted on glass slides.

Imaging was performed on a spinning disk confocal microscope (a Nikon Ti2 with a Yokogawa CSU-W1 spinning disk head, described(Zheng, 2024 #335). Z-stack images at 0.5-μm spacing were collected with a 60×/1.40 NA Plan Apochromat oil immersion objective (Nikon).

### Live-cell imaging

Live cell imaging (LCI) videos were acquired and analyzed using NIS-Elements (Nikon). To perform LCI, we used cells expressing both mCherry-H2B and GFP-Tubulin to capture mitotic events. LCI was performed on a Ti2 inverted microscope fitted with a CSU-W1 spinning disk system (Nikon). To capture mitotic cells, z-stacks (+6μm above and -4 μm below the focus plane at 0.5-μm spacing) were captured every 2 minutes for 2-4 hours using a Zyla 4.2 sCMOS camera, and a 20×/0.95 NA Plan Apochromat Lambda objective with the correction collar set to 0.17. For higher resolution imaging of cells undergoing mitosis, imaging was performed on a Nikon Ti2 inverted microscope fitted with a Yokogawa CSU-W1 spinning disk head. Z-stacks (+6 μm above and -4 μm below the focus plane at 0.75-μm spacing) were collected every 2 minutes for 2 hours, using Zyla 4.2 sCMOS camera, and a 100×/1.45 NA Plan Apochromat Lambda oil immersion objective (Nikon). An environmental enclosure was used to maintain cell culture conditions (37°C and humidified 5% CO2) for all live-cell confocal imaging. The timing of mitosis was measured from the point of DNA condensation to onset of anaphase. The spindle geometry was determined in metaphase cells prior to the onset of anaphase.

### Protein extraction and immunoblotting

Protein extracts were prepared in NETN buffer (50mM Tris 7.5, 1mM EDTA, 0.5% NP-40, 5% glycerol, 150mM NaCl, complete TM EDTA-free protease inhibitor cocktail (Sigma, 1183617001), and PhosSTOP^TM^ phosphatase inhibitor cocktail (Sigma, 04906837001)). Briefly, cells were harvested and washed three times in ice cold PBS before resuspending in lysis buffer. After 20 min incubation with end-over-end rotation at 4°C, the cell lysate was clarified by centrifuging at 13,000 rpm for 10 min. Protein concentration in clarified lysate was determined by Pierce BCA Protein Assay Kit (Thermo Scientific, 23225) with reference to a standard curve generated with BSA. Extracts were mixed with 4X Laemmli sample buffer and heated at 95 °C for 5 min. Denatured extracts were resolved on pre-cast NuPAGE 4–12%, 1.5mm Bis-Tris polyacrylamide gels (Life Technologies, NP0335) and transferred to 0.2 µm nitrocellulose membranes (BioRad, 1620112). Membranes were blocked in 5% BSA (Sigma, 10735086001) in TBS-1% Tween20 for 1 h and incubated in primary antibodies at 4°C for 16-18 hours. Membranes were washed three times the next day with TBS-0.05% Tween20 at room temperature with rocking. Membranes were then probed with HRP-conjugated goat anti-mouse (Jackson ImmunoResearch, 115035008) or goat anti-rabbit (Jackson ImmunoResearch, 111035008) for 1 hour (1:3000). Amersham ECL Western Blotting Detection Reagent Kit (GE Healthcare, RPN2106) was applied to develop the blots. For PLK1 and pT210 immunoprecipitation, anti-PLK1 or anti-pT210 antibodies were cross-linked to Protein G Dynabeads (Thermo Fisher Scientific #10004D). Cells were collected, washed with PBS, and lysed in lysis buffer [50 mM tris-HCl (pH 7.5), 150 mM NaCl, 1 mM MgCl2, 1 mM EDTA, 0.5% Triton X-100, 1 mM β-glycerophosphate, 1 mM sodium molybdate, 1 mM sodium fluoride, 1 mM sodium orthovanadate, and protease inhibitors]. The lysate was clarified by centrifugation at 13,000g for 30 min at 4°C. The supernatant was transferred to a new tube and incubated with antibody-coupled beads for 2 hours at 4°C with rotation. Afterward, beads were washed and eluted with 4X Laemmli sample buffer and heated at 95 °C for 10 min. Denatured extracts were resolved as mentioned above.

### Mass spectrometry

Samples were subjected to on-bead digestion of immunoprecipitated proteins, bead mixtures were subjected to cysteine reduction (1.5 mM dithiothreitol for 15 min at 30°C), followed by alkylation (7.5 mM iodoacetamide, 30 min at RT). Bead mixture was diluted to 2M Urea with 100 mM ammonium bicarbonate and digested with 100 ng trypsin (Promega) overnight at 37°C. After bead removal, peptides were acidified with 0.1% formic acid, extracted by solid phase extraction with C18 cartridges (Gracepure SPE C18-Aq), and dried by SpeedVac (Thermofischer). Lastly, extracted peptides were resuspended in 0.1% formic acid discovery mass spectrometric analyses were performed on a quadrupole orbitrap mass spectrometer equipped with a nano-electrospray ion source (Orbitrap Exploris 480, Thermo Scientific). Chromatographic separation of the peptides was performed by liquid chromatography (LC) on a Evosep system (Evosep One, Evosep) using a nano-LC column (EV1137 Performance column 15 cm x 150 µm, 1.5 µm, Evosep; buffer A: 0.1% v/v formic acid, dissolved in milliQ-H2O, buffer B: 0.1% v/v formic acid, dissolved in acetonitrile). Twenty µL of the diluted digests were injected in triplicate and separated using the 30SPD workflow (Evosep). The mass spectrometer was operated in positive ion mode and data-independent acquisition mode (DIA) using isolation windows of 16 m/z with a precursor mass range of 400-1000, switching the FAIMS between CV-45V and -60V with three scheduled MS1 scans during each screening of the precursor mass range. Acquired spectra were analyzed using Spectronaut 17.6.230428 (Biognosys) with the standard settings of the directDIA workflow except that quantification was performed on MS1 with a human SwissProt database (www.uniprot.org, UP000005640 release 2023_1, 20422 entries). For the quantification, the Q-value filtering was set to the classic setting with global normalization and no imputation. Normalized intensity values were processed using Perseus v1.6.15 (MaxQuant). Data was log2 transformed, after which the three technical replicate measurements per sample were grouped and filtered to keep each protein with three valid values in at least one group. Lastly, missing data was imputed with random values obtained from a gaussian distribution with a downshift of 1.8x standard deviation of each sample and a width of 0.3x the standard deviation in Perseus.

### Antibodies

The antibodies used in this study were as follows: mouse α-tubulin (B-7) Santa Cruz sc-5268; mouse β-actin (C4) Santa Cruz sc-47778; rabbit 53BP1 Thermo-Fisher Scientific #NB100304; mouse 53BP1 Thermo Fisher Scientific) #MAB3802MI; rabbit TIRR Sigma Aldrich#HPA04418; mouse PLK1 antibody [35-206] Abcam # ab17056; mouse PLK1 (phospho T210) antibody [2A3] Abcam # ab39068; CENPF Abcam # ab5-50ul; H4K20ME2 PAB Life Technologies # 720085; NDC80 (9G3.23) Thermo Fisher Scientific # NB100338; CENPT Abcam # ab86595; CENP-A Thermo Fisher Scientific #PA5-17194; CENPB Abcam # **ab25734; BubR1** Abcam #**ab70544;** CENPC Abcam #**ab50974**; MIS18 milipore HPA020585; phospho-MIS18BP1 (Thr702) milipore #384187; and human anti-centromere Centromere Antibodies Incorporated 15–235.

### siRNAs

ON-TARGETplus Human TP53BP1 (7158) siRNA-SMARTpool, Horizon Discovery # L-003548-00-0005; ON-TARGETplus Human PLK1 (5347) siRNA Horizon Discovery # L-003290-00-0005; ON-TARGETplus Human NUDT16L1 (84309) siRNA - SMARTpool, Horizon Discovery # L-014872-02-0005;

### Spindle geometry, inter-kinetochore distance, kinetochore-microtubule (k-MT) attachment and cold shock assays

RPE-1 cells were treated with 9µM of RO-3306 for 16-18 hours to arrest cells at G2/M border. 16-18 hours post initial RO-3306 treatment, the cells were released from G2/M arrest by washing with pre-warmed media four times and addition of fresh media containing 20µM proTAME for the spindle geometry and IKD experiments. The cells were then fixed with 3.5% of PFA and immunostained as described above. For cold shock assay, the cells were released from G2/M arrest by washing with pre-warmed media four times into fresh media containing 5µM MG-132 for 2 hours. The cell culture dish was then placed on ice for 30 mins. The cells were then fixed with cold methanol for 10 minutes on ice and immunostained as described above. Inter-kinetochore distance and cold shock assay were performed and analyzed as previously described^62^. Attachment of kinetochores to microtubules was assessed by examination of the full series of Z-axis focal planes. Kinetochores in regions where fibers were not easily visualized were not included in the analysis.

