## Supplementary figures and images for "The TIRR–53BP1 axis controls PLK1 spatiotemporal regulation during mitosis" (from another version of this preprint: version 1, 2026-06-30)

### Supplemental Figures

#### Supp figure 1

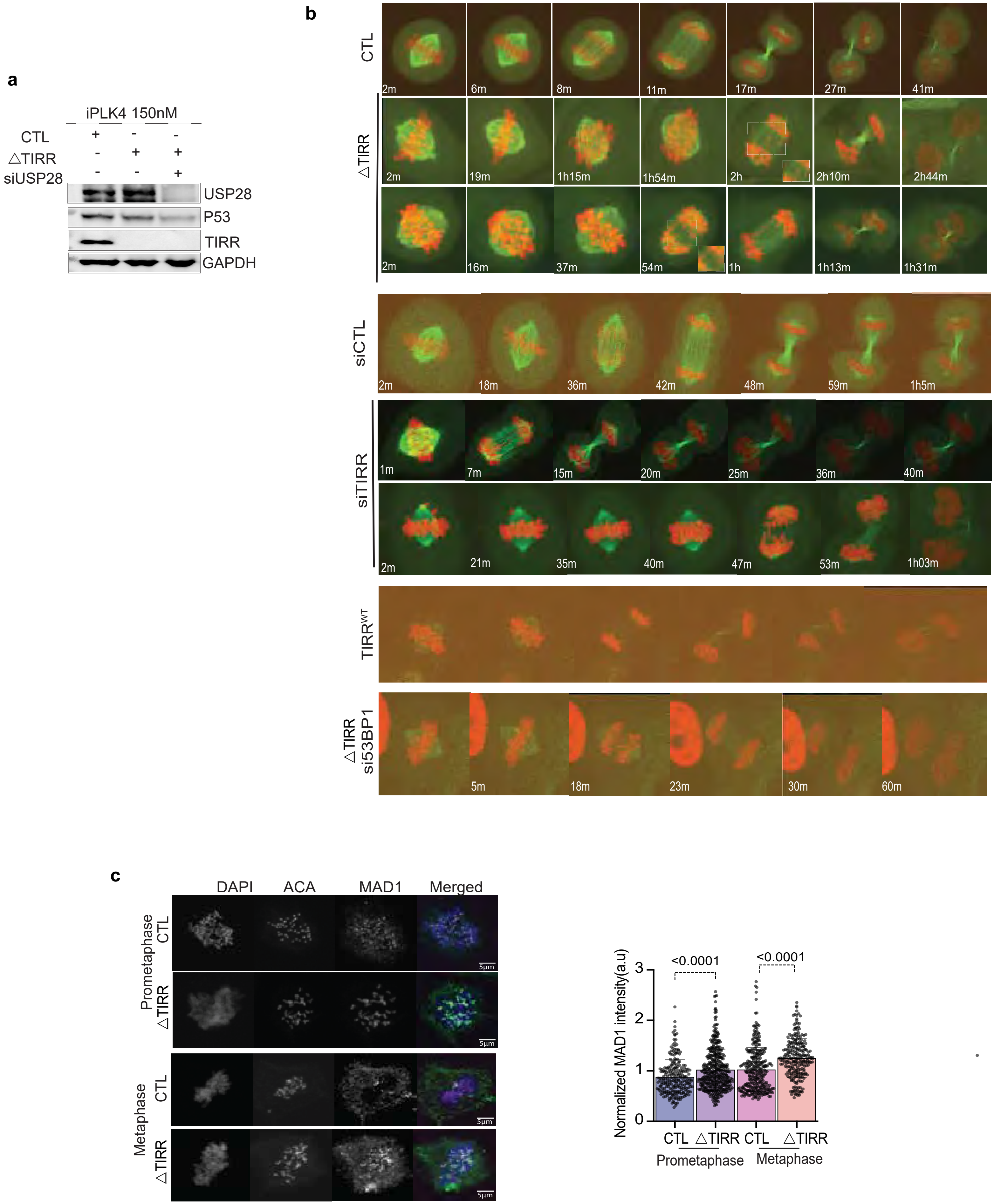

#### Supp figure 2

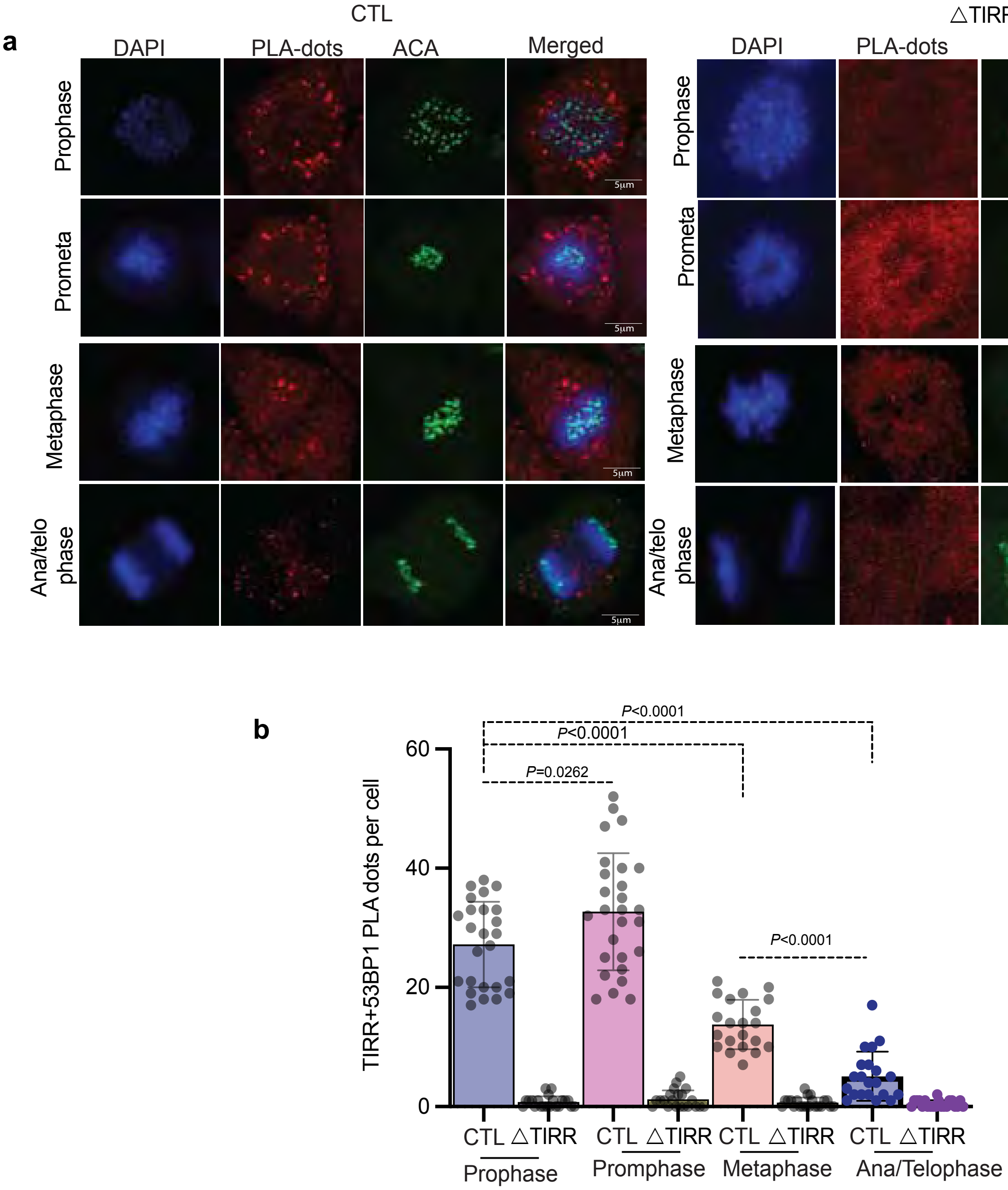

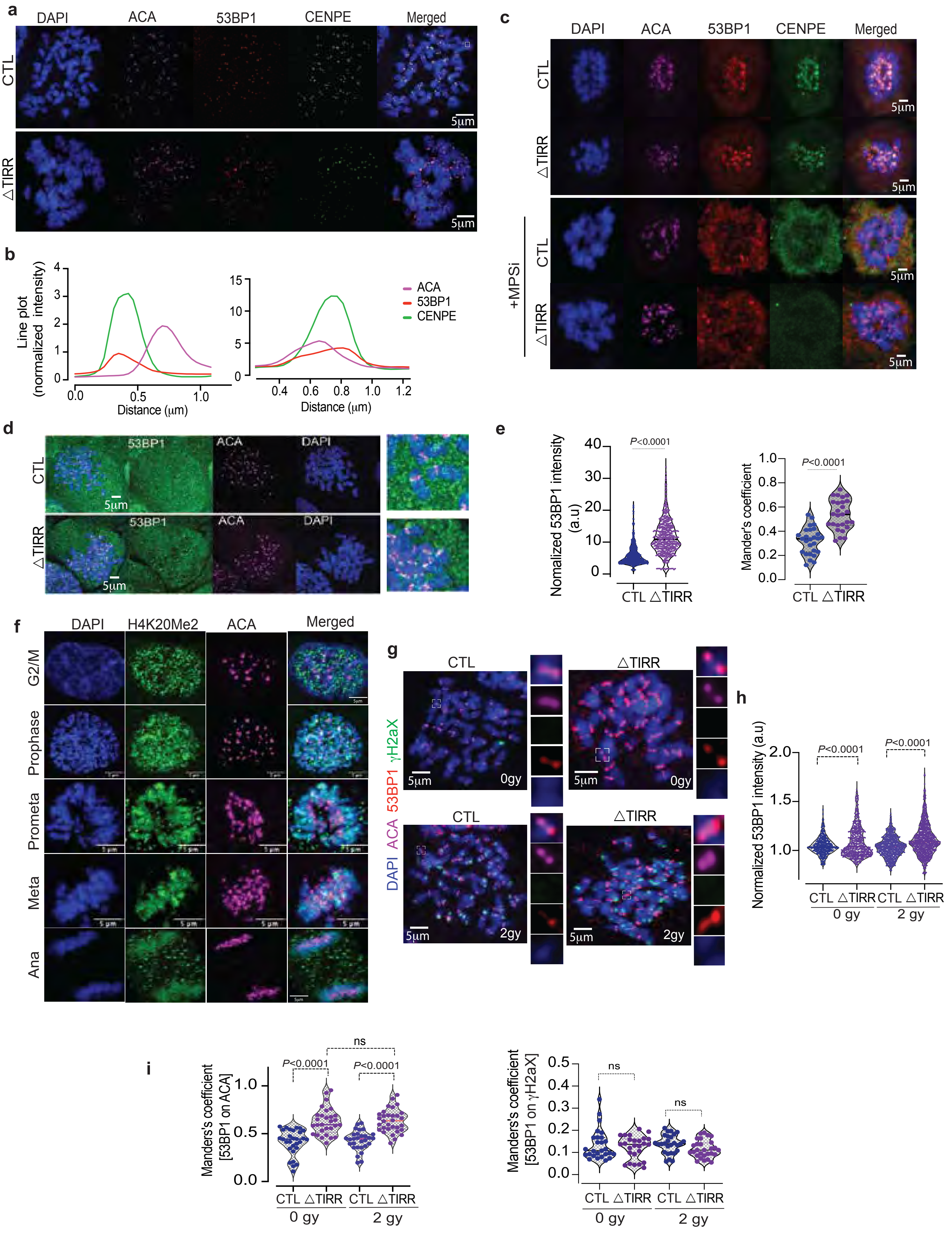

**Supp figure 4**

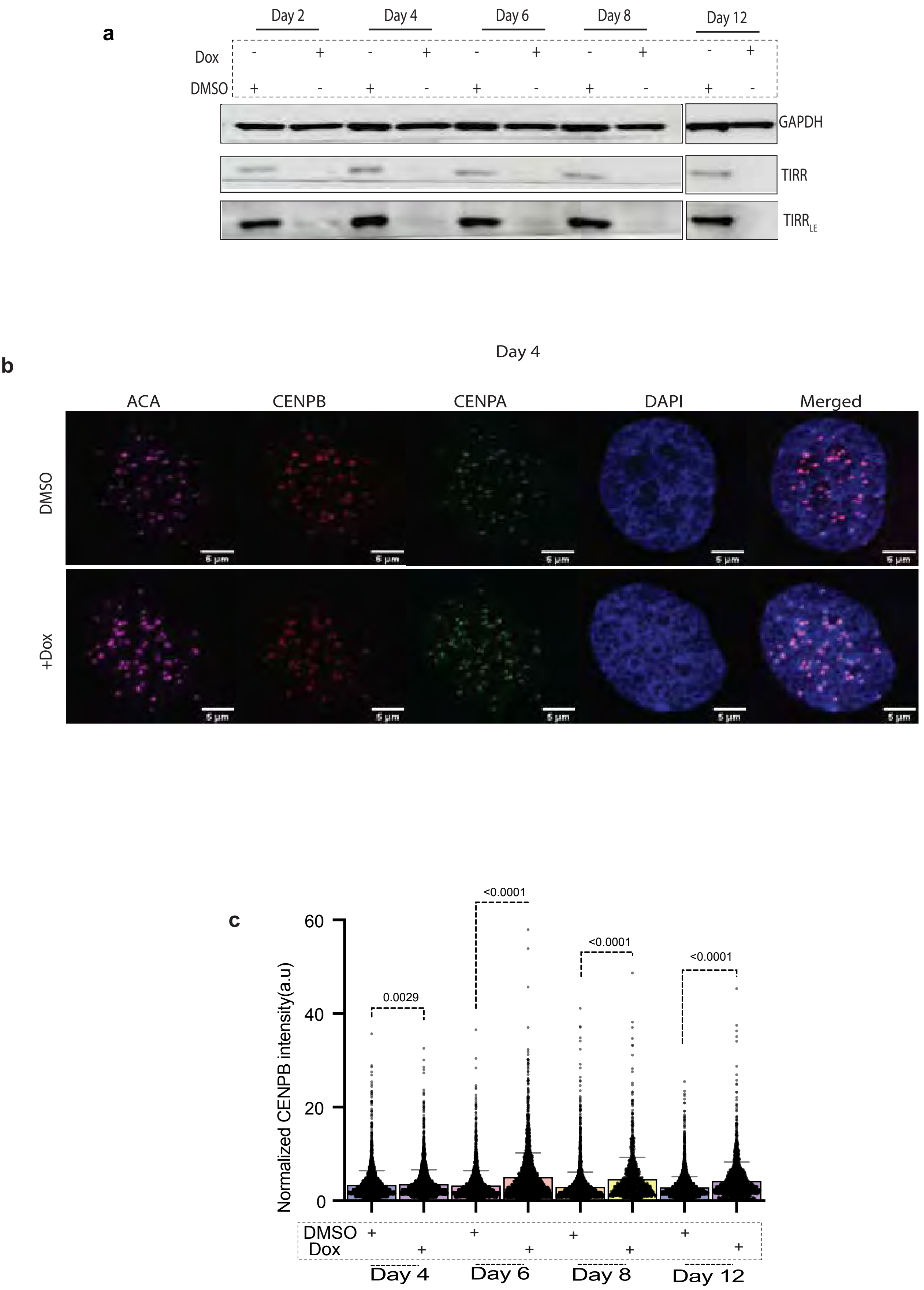

**Supp figure 5**

**a**

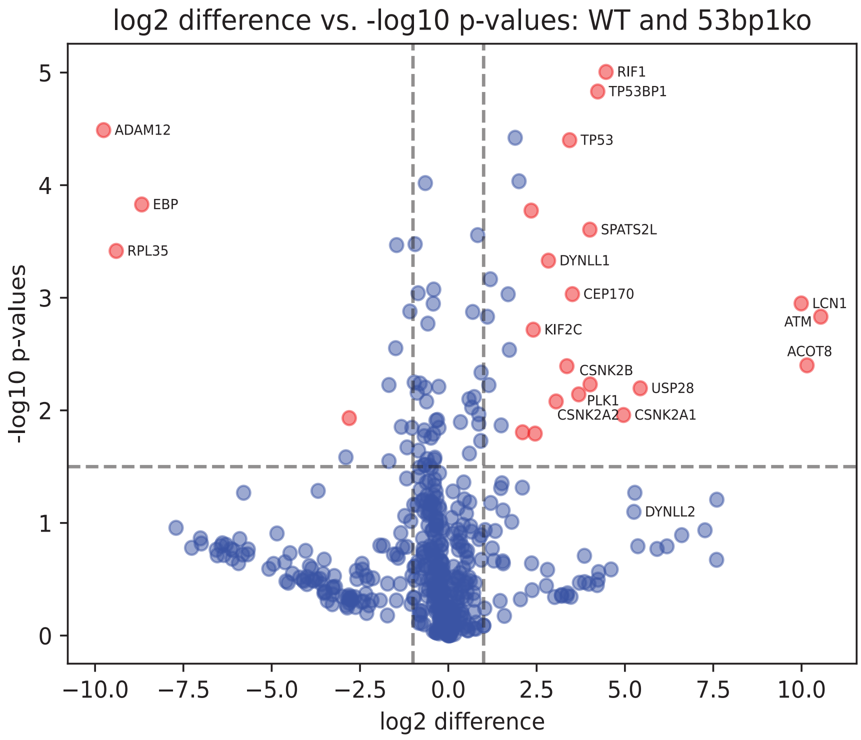


**b**

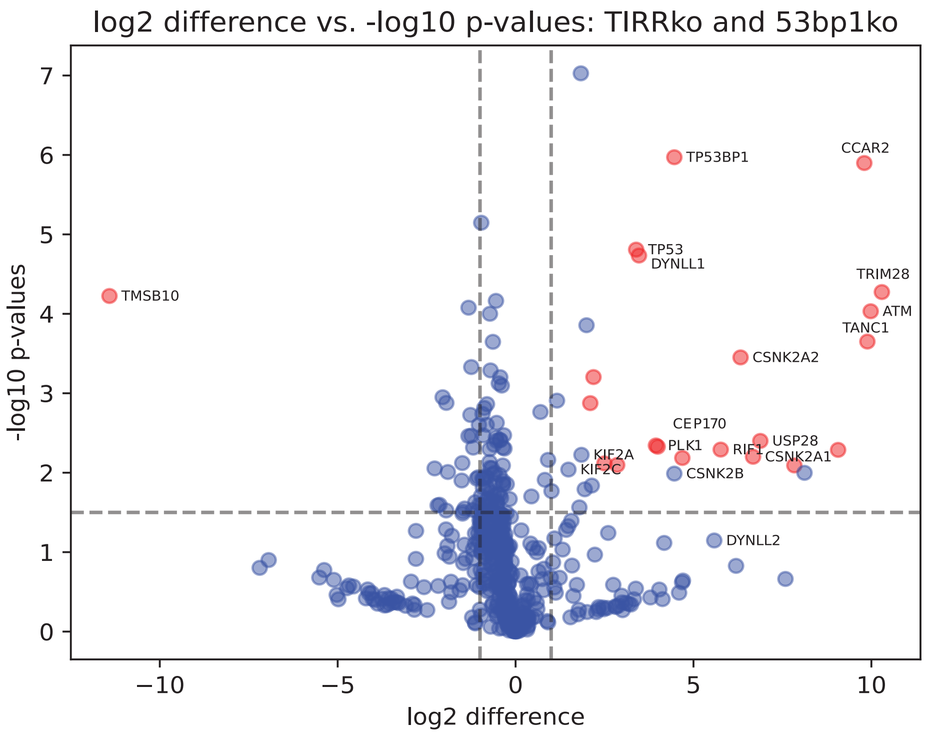
